# Temporal evolution of hemodynamics in murine arteriovenous fistula: a micro-CT based computational fluid dynamics study

**DOI:** 10.1101/2025.03.24.644912

**Authors:** Lianxia Li, Unimunkh Uriyanghai, Christine Wai, Hong Yuan, Eric W. Livingston, Edward M. Bahnson, Vinay Sudarsanam, Samuel Haddad, Prabir Roy-Chaudhury, Boyce E. Griffith, Gang Xi

## Abstract

In this study, we investigated the hemodynamic characteristics of arteriovenous fistulae (AVF) in murine models using micro-CT based computational fluid dynamics (CFD). By combining high-resolution micro-CT imaging with ultrasound flow measurements, our methodology offers a cost-effective and efficient alternative to traditional MRI-based approaches. CFD simulations performed at 7 and 21 days post-surgery revealed significant temporal changes in both geometry and hemodynamics. Geometric analysis showed that: the proximal artery diameter increased from 0.29 mm to 0.38 mm, while the initial 2 mm fistula segment showed a 21.6% decrease (0.74 mm to 0.58 mm). Blood flow through the AVF nearly doubled from 1.33 mL/min to 2.57 mL/min. Time-averaged wall shear stress (TAWSS) peak values increased from 142 Pa (day 7) within the proximal artery to 200 Pa (day 21), in the stenotic region. The oscillatory shear index (OSI) showed marked elevation at the anastomosis (increasing from 0.22 to 0.48), indicating disturbed flow development. An inverse relationship between TAWSS and OSI was identified consistent with previous studies. Our methodology demonstrates the capability to analyze relationships between early hemodynamics and subsequent geometric changes. This approach could enable identification of regions susceptible to stenosis development and monitoring of AVF maturation, which could ultimately lead to quantitative metrics to evaluate surgical outcomes and early therapeutic interventions.

## Introduction

Arteriovenous fistulae (AVF) are surgically created connections between an artery and a vein, primarily used for hemodialysis access in patients with end-stage renal disease (ESRD) [1]. Despite being the preferred vascular access method due to superior longevity and lower complication rates compared to central venous catheters and grafts [2], AVF maturation and failure remain significant clinical challenges [3]. Maturation failure, defined as an inability of the AVF to increase flow and diameter adequately to support a successful clinical dialysis session occurs in approximately 20-50% of cases, leading to increased morbidity and healthcare costs [4–6].

Understanding the hemodynamic environment within the AVF is crucial, as it plays a pivotal role in both the maturation process and long-term patency [5, 7], with factors such as blood flow velocity, pressure, and wall shear stress significantly influencing endothelial cell function, vascular remodeling, and the development of intimal hyperplasia, a leading cause of AVF failure [7]. The creation of an AVF also triggers cardiac changes in murine models [8], suggesting a complex interplay between cardiac function and local hemodynamics. Previous studies suggest that stenosis and neointimal hyperplasia development are associated with low wall shear stress but high oscillatory shear index (OSI) [9] [10], while higher wall shear stress with a lower OSI promote lumen expansion after AVF creation [11]. However, the complete mechanistic relationships between these hemodynamic parameters and vascular remodeling are complex and have not been fully elucidated [12, 13]. Further, traditional experimental methods for studying AVF hemodynamics have been limited by insufficient spatial and temporal resolution to capture complex flow dynamics [14–16].

Computational fluid dynamics (CFD) offers a powerful tool to overcome these limitations by enabling detailed simulations of blood flow within AVF [17]. By integrating anatomical data from high-resolution imaging modalities such as micro-computed tomography (micro-CT), CFD provides comprehensive insights into the hemodynamic environment within different regions of the AVF. This approach enables precise quantification of flow patterns, pressure distributions, and wall shear stress profiles, facilitating a deeper understanding of factors contributing to AVF maturation and failure [18].

Murine models serve as valuable platforms for AVF research because of their genetic similarities to humans, the availability of sophisticated genetic tools, the ability to manipulate individual genes and their downstream proteins, and cost-effectiveness [19, 20]. While MRI-based CFD models have been developed for murine AVF analysis [21], CT-based CFD models with ultrasound velocity data offer several distinct advantages that have not been fully explored. For geometric reconstruction, in vivo CT imaging provides not only high spatial resolution, but accurate vasculature morphology under live physiological or pathophysiological condition, which is crucial for more precise computational modeling of complex AVF geometries [22, 23]. For flow measurements, Doppler-based ultrasound methods offer significant advantages over phase-contrast MRI-based flow measurement. Although MRI-based flow measurement is usually more accurate and provides 3D flow, Doppler ultrasound enables quick, repeated assessments with real-time feedback that allows for a detailed profile of CFD changes over time, in addition to high portability, and cost-effectiveness [24].

Thus, our approach combines the strengths of both modalities: high-resolution CT imaging for precise geometric reconstruction and Doppler ultrasound for efficient flow measurements. This combination also offers advantages over 4D flow MRI in terms of spatial resolution and reduced scan times [25, 26], which is particularly beneficial for imaging small vessels near bony structures such as the neck region where AVFs are typically created.

While previous studies have characterized either the geometric or hemodynamic aspects of AVF development[16, 27, 28], the relationship between early flow patterns and subsequent vascular remodeling remains poorly understood. This study aims to bridge this gap by:

(1) Developing a micro-CT-based CFD methodology for murine AVF analysis.
(2) Characterizing the temporal evolution of AVF hemodynamics with high spatial resolution.

By integrating detailed anatomical data from high-resolution imaging with advanced computational techniques, we seek to establish quantitative metrics for predicting AVF maturation outcomes. Our findings are expected to provide valuable insights into the mechanistic underpinnings of AVF development, potentially leading to future clinical interventions and improved outcomes for hemodialysis patients in the future.

## Materials and methods

### Animal model

Male C57BL/6J mice, aged 14-16 weeks obtained from Jackson lab, were used in this study. Imaging and ultrasound data were collected at days 7 and 21 post-surgery to characterize the early and late stages of AVF development.

#### Ethical considerations

All animal procedures were approved by institutional guidelines and approved by the University of North Carolina at Chapel Hill at the Institutional Animal Care and Use Committee (IACUC) and performed in accordance with National Institute of Health guidelines. Proper care was taken to minimize animal suffering.

#### Surgical procedure

AVF surgery was performed following established protocols. The mouse was anesthetized using a combination of isoflurane inhalation (1-3%) with prior surgery analgesia (buprenorphine, 0.1 mg/kg.

A 1 cm longitudinal incision was made in the middle of the neck. Under a surgical microscope (12.5-25X magnification), the right external jugular vein and common carotid artery were carefully exposed through blunt dissection. The jugular vein was ligated distally and divided, while the carotid artery was temporarily clamped using micro-vascular clamps. An end-to-side anastomosis was created between the jugular vein and carotid artery: first, an arteriotomy (approximately 0.4-0.5 mm) was made in the carotid artery, then the jugular vein was anastomosed to the artery using interrupted 10-0 nylon sutures. Fig 1 illustrates the surgical creation of the AVF, showing the end-to-side anastomosis between the jugular vein and carotid artery.

**Fig 1.**
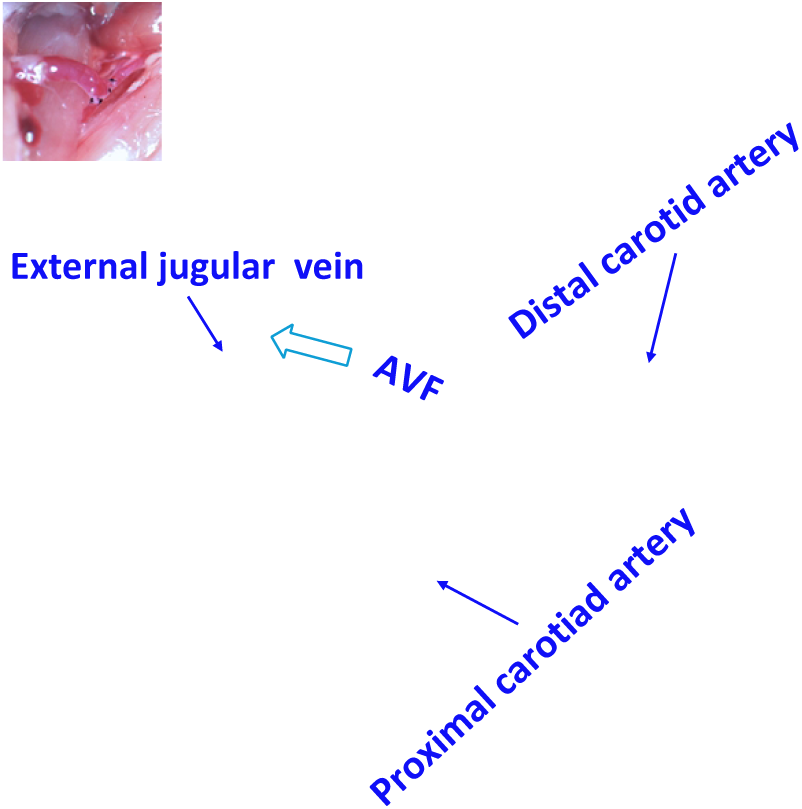
AVF creation using end-to-side anastomosis between jugular vein and carotid artery in murine model. White arrow shows direction of venous outflow.

#### Imaging protocol

High-resolution micro-computed tomography (micro-CT) was performed in vivo at days 7 and 21 post-surgery using a Quantum GX2 system (Revvity; Waltham, MA USA) with the following parameters: 90 kV of peak energy, 88 μA of current, 72 μm isotropic voxel size, and 2 min of acquisition time. For day 21 imaging, an additional scan with 14 min acquisition time was performed in order to achieve a high signal-to-noise ratio. Prior to imaging, either Fenestra-HDVC (day 7) or mvivoAU (day 21) (MediLumine Inc, Montreal, Quebec CA) was administered intravenously via tail vein catheter (5 µl/g for both agents) to provide vascular contrast for vessel segmentation.

#### Blood flow velocity acquisition

A high-frequency Doppler ultrasound system (Vevo F2, FUJIFILM VisualSonics) with a UHF57x linear array transducer with center frequency at 32 MHz was used to obtain blood flow velocity measurements on the same day, prior to micro-CT imaging. Mice were anesthetized using 1.5-2% isoflurane, and body temperature was maintained at 37°C during measurements.

Velocity measurements were obtained using PW Doppler mode at three locations: the proximal artery (1.5 mm upstream of anastomosis), distal artery (1.5 mm downstream of anastomosis), and the venous segment of the AVF. The angle between the wave beam and flow direction was kept below 60 degree either by animal positioning or by using additional 15° beam steering. At each location, pulsatile velocity waveforms were recorded over multiple cardiac cycles. Three consecutive cycles were extracted and averaged to generate representative velocity profiles for each location. Peak velocity in mm/s was determined for each location. The volume flow rate was calculated using Q = AV, in which A is the cross-sectional area obtained from the micro-CT data, and V is the mean velocity. The cross-sectional velocity was assumed to follow a parabolic profile, with the average velocity being half of the maximum velocity measured at the centerline.

Heart rate was monitored throughout the measurements and remained stable at approximately 450 beats per minute. The acquired velocity data served as boundary conditions for the subsequent CFD simulations.

### Image processing and geometry reconstruction

Image processing and three-dimensional vessel reconstruction were performed using a multi-step approach. The micro-CT scans were first processed in ITK-SNAP (Version 4.2) [29, 30] for initial segmentation (Fig 2(a)).

**Fig 2.**
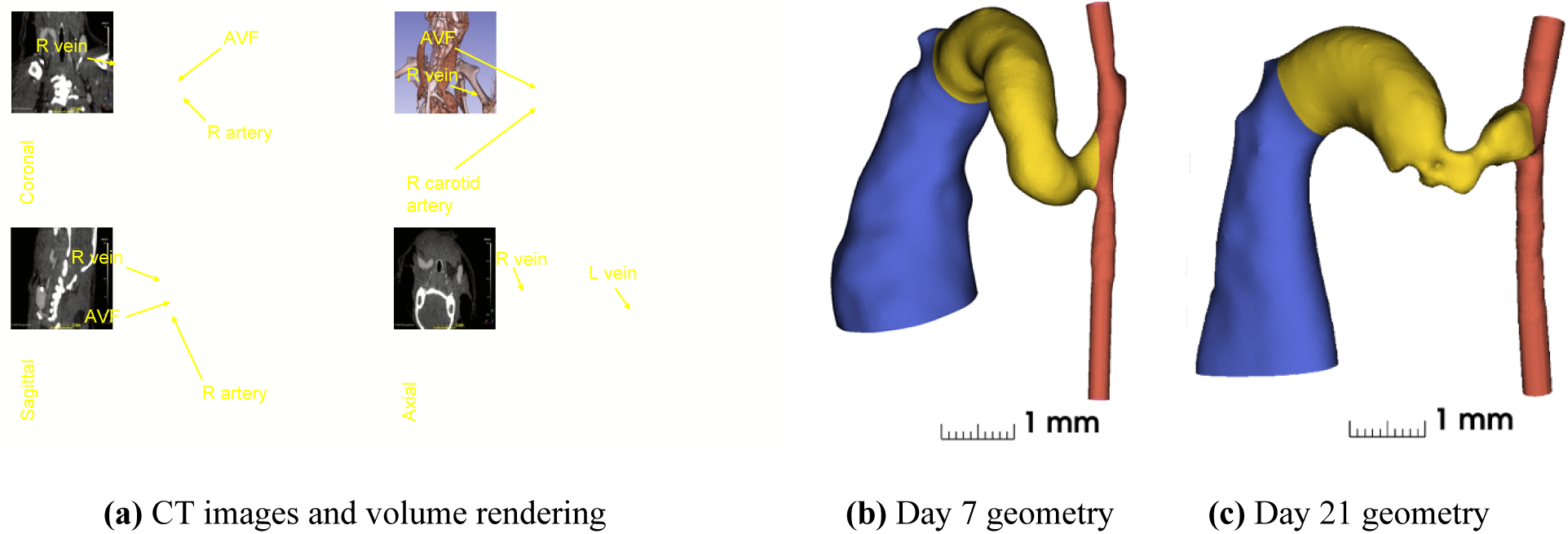
(a) CT images and volume rending for day 7 in Slicer 3d [33]; three-dimensional reconstructions of murine arteriovenous fistula (AVF) models showing vascular architecture at (b) day 7 and (c) day 21 post-surgery. Color coding represents distinct vessel segments: red indicates the carotid artery (with the proximal segment at bottom and distal segment at top), yellow highlights the AVF region (4 mm of length), and blue shows the main venous drainage with its characteristic ballooning of the vein starting 4 mm from the anastomosis.

The vascular lumen was segmented using a combination of threshold-based and region-growing techniques. Initial thresholding (300-1000 HU) isolated contrast-enhanced vessels and bones from the neck area, followed by seed-based region-growing to further segment connected voxels to form vessel lumens. Manual editing was performed where necessary, particularly at the anastomosis region to ensure accurate geometry capture.

The segmented vasculature was exported as STL files and further processed in Autodesk Meshmixer [31] to remove artifacts and smooth surface irregularities. Inlet and outlet regions of the artery were extended by 2 mm to ensure fully developed flow, and geometric landmarks were defined for consistent analysis.

Centerline analysis was performed using VMTK (the Vascular Modeling Toolkit [32]) to extract vessel centerlines and calculate cross-sectional areas and diameters. These measurements provided geometric metrics for quantitative comparison between timepoints. The final reconstructed geometry consisted of the proximal artery (inflow), distal artery (outflow), and the arteriovenous fistula, which was defined as the 4 mm length of vessel extending from the anastomosis as shown in Fig 2(b) and (c).

### Computational fluid dynamics

The simulation was conducted using OpenFOAM v2306 [34], an open-source finite volume based computational fluid dynamics software package. OpenFOAM is written in C++ and provides numerical solvers, utilities, and tools for solving continuum mechanics problems involving fluid flow, chemical reactions, and heat transfer, among others [35].

#### Governing equations

The blood flow within the arteriovenous fistula (AVF) is modeled using the incompressible Navier-Stokes equations:

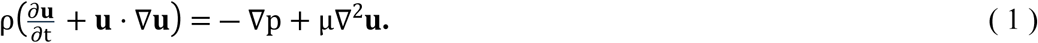

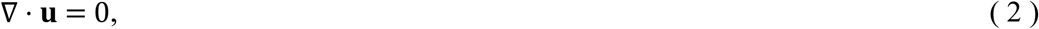

in which ρ is the density of blood (1,040 kg/m³), μ is the dynamic viscosity of blood (4 × 10^―3^ Pa ⋅ s), **u** is the velocity vector of the blood flow, p is the pressure, and t is the time.

#### Mesh generation and grid independence

The computational domain encompasses the proximal artery, distal artery, fistula (4 mm length from anastomosis) (Fig 3), and vein. The domain was discretized using tetrahedral meshes with five prismatic boundary layers (first layer thickness 1-3 µm, growth ratio 1.2) near the walls for accurate flow gradient calculations. Grid independence was established using three mesh resolutions for the day 7 model: coarse (0.62M elements), medium (1.34M elements), and fine (2.52M elements). With differences in velocity and pressure distributions between consecutive refinements below 1%, the medium mesh resolution was adopted for all analyses, yielding 1.34M and 1.97M elements for day 7 and day 21 models, respectively.

**Fig 3.**
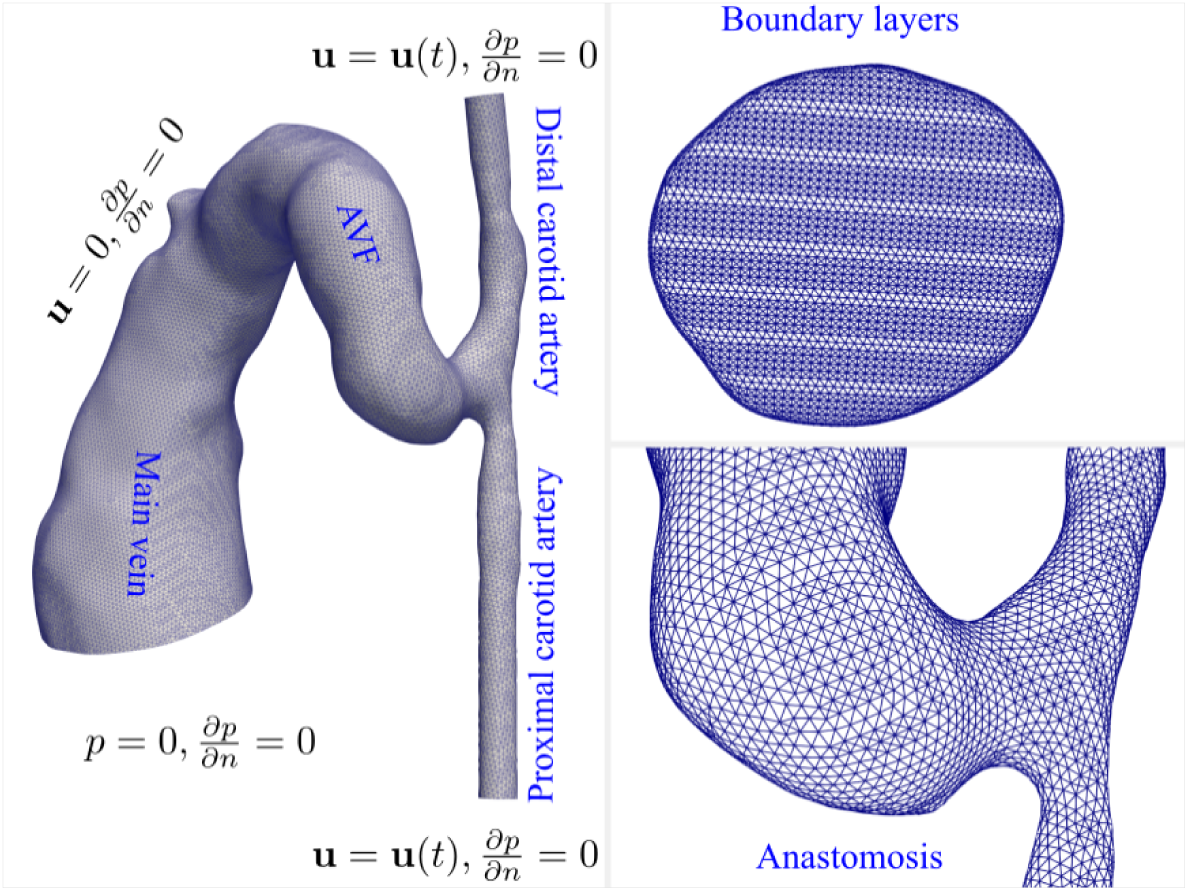
Computational domain, boundary conditions, and mesh configuration for the murine AVF model (day 7 shown). Boundary conditions are specified as follows: (1) velocity-based inlet conditions at both proximal and distal carotid artery ends, derived from ultrasound Doppler waveforms; (2) no-slip conditions at all vessel walls; and (3) pressure boundary condition at the venous outlet. The computational domain is discretized using tetrahedral meshes with five boundary layers near the walls. The inset shows a detailed view of the mesh structure at the arteriovenous junction.

#### Boundary conditions and numerical methods

Velocity boundary conditions from ultrasound Doppler measurements were applied at artery inlets, with zero-pressure condition at the venous outlet (all pressures reported relative to outlet). No-slip conditions were enforced at vessel walls. Time discretization used backward Euler scheme, while spatial discretization employed second-order upwind schemes with Gauss linear gradient calculations. The SIMPLE algorithm handled pressure-velocity coupling, with 10⁻⁶ convergence criteria. Simulations ran for three cardiac cycles (CFL = 0.1), with the final cycle used for analysis.

### Data analysis methods

#### Hemodynamic parameters

Several hemodynamic parameters were calculated to characterize the flow field and its potential influence on vascular remodeling. The wall shear stress (WSS), representing the tangential force exerted by blood flow on the vessel wall, was computed as

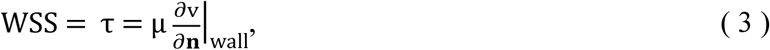

in which v is the velocity component parallel the vessel wall and **n** is the unit vector normal to the wall.

The time-averaged wall shear stress (TAWSS) is calculated over one cardiac cycle (T)

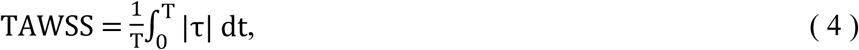

TAWSS provides a measure of the chronic mechanical load experienced by endothelial cells [7]. In AVF development, regions of consistently high TAWSS often trigger outward vascular remodeling as an adaptive response, whereas regions of low TAWSS may be prone to intimal hyperplasia development [21].

The oscillatory shear index (OSI) quantifies the directional changes in wall shear stress over the cardiac cycle

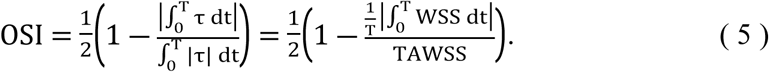

OSI ranges from 0 to 0.5, where 0 indicates unidirectional flow and 0.5 represents complete flow reversal. In AVF, high OSI regions often correlate with disturbed flow patterns and are frequently associated with stenosis development [36]. This is particularly significant at the anastomosis and in post-stenotic regions where complex flow patterns emerge [23].

The vortex structure is commonly discerned using the Q-criterion, computed as the second invariant of the velocity gradient tensor:

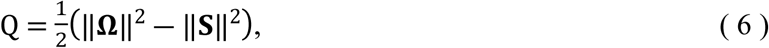

in which **Ω** and **S** are the antisymmetric and symmetric components of the velocity gradient tensor, respectively. Positive Q-values indicate regions where rotation dominates strain, identifying coherent vortical structures [22].

#### Data analysis

Cross-sectional averages and maxima of hemodynamic parameters were computed at key locations along the vessel centerline. Temporal comparisons between day 7 and day 21 were performed for both geometric and hemodynamic parameters. Correlations between early hemodynamic conditions and subsequent geometric changes were analyzed to identify potential predictive relationships.

## Results and discussion

### Clinical analysis

#### Temporal changes in vessel geometry

Table 1 summarizes the lumen diameters of the carotid artery, fistula, and vein. For each segment, the values in parentheses indicate the minimum and maximum diameters. Additionally, detailed statistical analyses of the AVF region are provided for two specific segments: the initial 2 mm segment adjacent to the anastomosis and the extended 4 mm segment, to characterize the spatial variation in remodeling patterns.

**Table 1.**
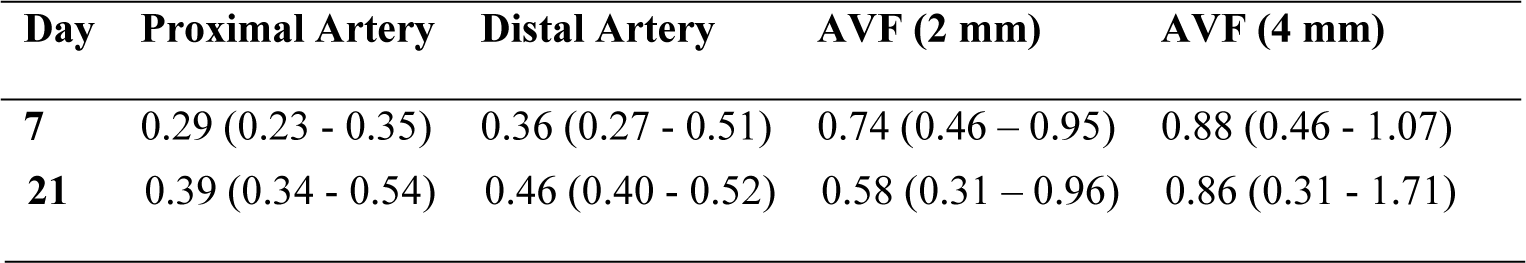
Diameters of the AVF models for day 7 and day 21.

Temporal analysis revealed distinct changes in vessel geometry between day 7 and day 21. The proximal carotid artery showed a uniform expansion, with the average diameter increasing from 0.29 mm to 0.38 mm. In contrast, the fistula demonstrated more complex remodeling patterns that varied by region. Analysis of different fistula segments revealed varying degrees of remodeling: while the average diameter over the initial 4 mm segment remained relatively stable (slight decrease from 0.88 mm to 0.86 mm), a more focused analysis of the initial 2 mm segment showed notable narrowing, with the average diameter decreasing by 21.6% from 0.74 mm to 0.58 mm. The most severe narrowing was observed at the point of minimum diameter, which decreased substantially from 0.46 mm to 0.31 mm (32.6%). The vein exhibited the most dramatic change in the opposite direction, with its maximum diameter expanding from 1.07 mm to 1.71 mm (59.8%).

Comparison of the longitudinal profiles of lumen diameter on day 7 and day 21 is demonstrated in Fig 4. Both timepoints show a characteristic narrowing downstream of the anastomosis, but with distinct patterns. On day 7, the narrow region was localized and positioned close to the anastomosis. By day 21, the vessel geometry had remodeled such that the immediate anastomosis region showed expansion, while the stenotic segment had both elongated and shifted further downstream. This temporal evolution of the stenosis pattern suggests dynamic vascular remodeling during the maturation process, which has been also observed in human AVFs [37].

**Fig 4.**
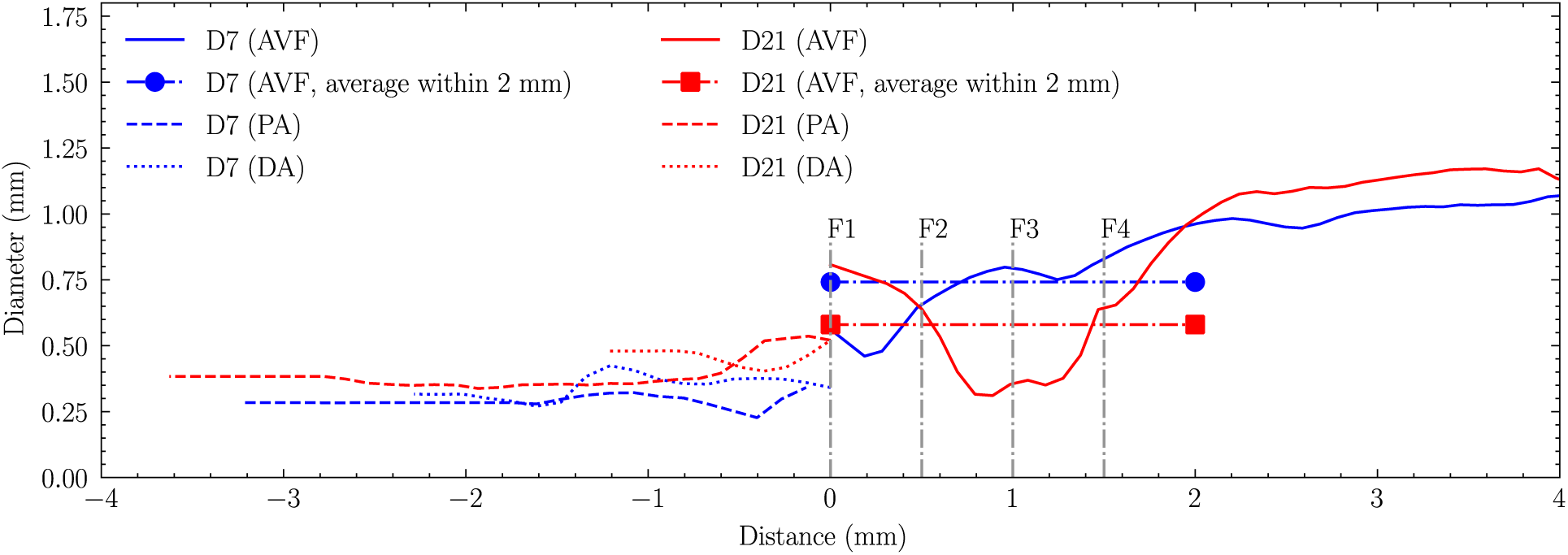
Lumen diameter profiles along the vascular segments at day 7 (D7) and day 21 (D21). The x-axis represents the distance from the anastomosis, with negative values indicating upstream segments (proximal artery [PA] and distal artery [DA]) and positive values indicating downstream segments (arteriovenous fistula [AVF]). The average AVF diameter is measured within the first 2mm of the fistula. Cross-sections F1-F4, spaced 0.5mm apart, indicate locations analyzed for hemodynamics in results.

#### Flow rates

Flow analysis revealed a significant temporal increase in total flow, with the AVF flow rate nearly doubling from 1.33 mL/min at day 7 to 2.57 mL/min at day 21, indicating fistula maturation. The flow distribution pattern demonstrated that the AVF served as the primary conduit at both timepoints, with 80% of proximal artery flow directed through the AVF at day 7, decreasing slightly to 70% by day 21, suggesting stenosis development within the AVF. Consequently, the proportion of blood flow through the distal artery increased from 20% to 30% between day 7 and day 21. Note that the blood in the distal artery flowed away from, rather than toward, the AVF junction. The proximal artery flow rate doubled over this period, while the heart rate maintained at approximately 450 beats per minute throughout the study period.

The typical carotid artery flow rate in normal mice ranges from 0.3 to 2.1 mL/min [38]. A 10 to 20-fold increase in blood flow rate is considered necessary for sufficient AVF maturation in humans [39]. Therefore, the flow rate increase observed in this study suggests that the AVF may not have achieved full maturation, possibly due to the development of stenosis.

### Hemodynamics analysis

#### Flow field characteristics

Flow field analysis revealed distinctive patterns at both timepoints, characterized by complex three-dimensional structures. The flow patterns illustrated in Fig 5 showed streamlines and velocity distributions at peak systole. Flow separation occurred at regions of high curvature and abrupt lumen changes, resulting in asymmetric velocity profiles at these locations.

**Fig 5.**
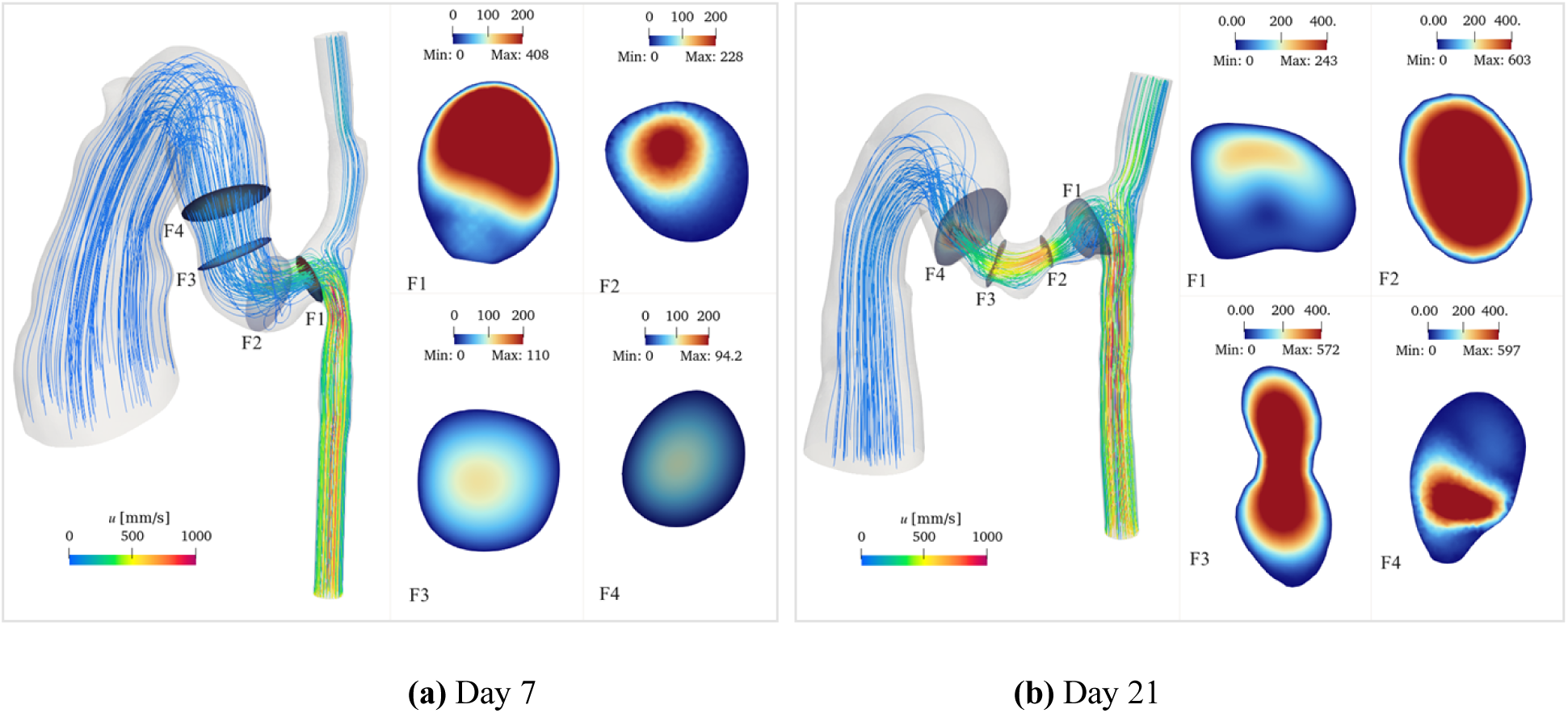
Flow patterns within the AVF at peak systole for day 7 (a) and day 21 (b) models. Streamlines colored by velocity magnitude show the three-dimensional flow trajectories. Cross-sectional velocity distributions at representative locations demonstrate the spatial evolution of the flow field. The color scale represents velocity magnitude in mm/s. Cross-sectional dimensions are not to scale.

Temporal and spatial comparison revealed distinct velocity patterns. At the anastomosis, the flow velocity was higher on day 7 (maximum velocity 408 mm/s) compared to day 21 (maximum velocity 243 mm/s), with pronounced asymmetry between the upper and lower regions. In contrast, at the stenotic region, day 21 exhibited substantially higher velocities (maximum 603 mm/s) compared to day 7 (maximum 228 mm/s), consistent with the increased flow constriction.

The flow symmetry showed different recovery patterns between the two timepoints. On day 7, the flow velocity profile became symmetric beyond the 2 mm fistula region. However, on day 21, flow asymmetry persisted beyond this region, as evidenced by both streamline patterns and cross-sectional velocity distributions. Further downstream, in the main vein, the flow exhibited axisymmetric patterns in cross-sectional views, with peak velocities concentrated along the vessel centerline, consistent with developed flow characteristics in straight vessel segments.

The centerline analysis of velocity and pressure (Fig 6) demonstrated characteristic flow features: velocity sharply increased before the stenosis, followed by deceleration, with peak velocities occurring within 1 mm downstream of the anastomosis. In accordance with the Bernoulli principle, an abrupt pressure drop was observed immediately preceding the stenotic region. Notably, pressure drop characteristics differed between timepoints: at day 21, 99% of the total pressure drop occurred within 1.5 mm downstream of the anastomosis, compared to only 50% at day 7 over the same distance.

**Fig 6.**
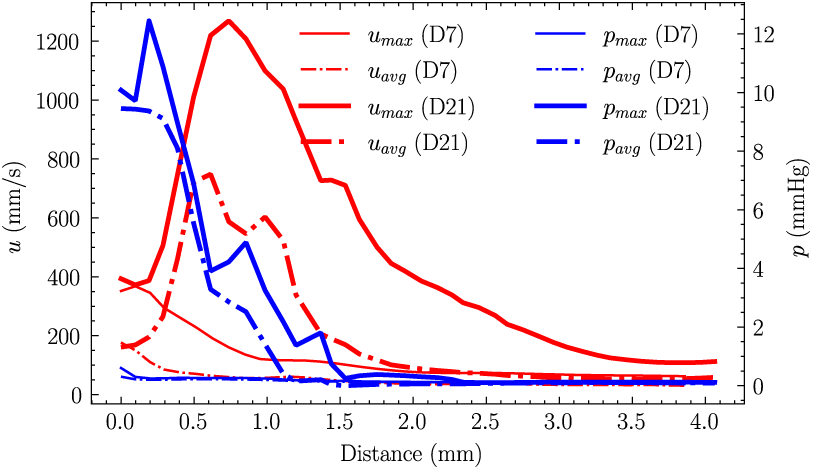
Cross-sectional average velocity and pressure along the vessel centerline at peak diastole for day 7 and day 21 models. The plots show velocity (red) and pressure (blue) distributions relative to distance from the anastomosis. Both models exhibit characteristic features: a sharp velocity increase preceding the stenosis followed by deceleration, with large velocities occurring within 1 mm downstream of the anastomosis. A corresponding abrupt pressure drop is observed immediately before the stenotic region. The pressure drop completes (99%) within 1.5mm downstream of the anastomosis on day 21, while it only occurs 50% over the same distance on day 7.

According to Poiseuille’s Law, the calculated flow resistance in the AVF segment (4mm length) on day 21 was approximately 9 times greater than that on day 7. This substantial increase in flow resistance indicates significantly higher friction forces, primarily attributable to the more severe stenosis developed by day 21.

#### Wall shear stress analysis

Fig 7 illustrates the temporal and spatial evolution of TAWSS along the AVF centerline. On day 7, TAWSS distribution shows distinct spatial patterns, reaching its peak within 0.5 mm downstream from the anastomosis, with the 4 mm fistula region showing an average value of 3 Pa. This localized peak suggests early flow adaptation near the surgical connection.

**Fig 7.**
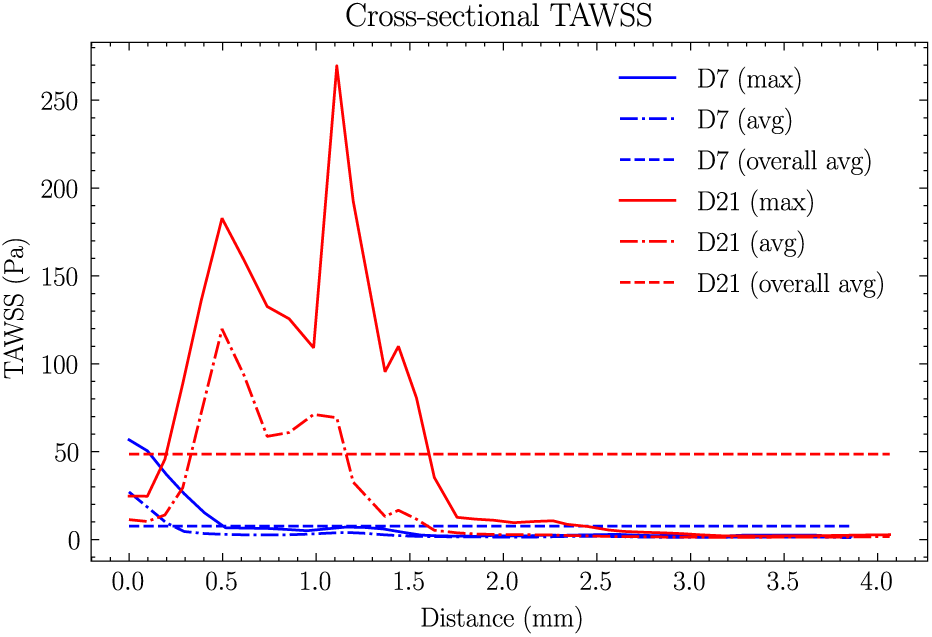
Cross-sectional maximum and average distributions of TAWSS along the centerline for (a) day 7 and (b) day 21. The two horizontal lines represent the average values across the first 4mm AVF. For day 7, peak TAWSS is localized within 0.5 mm downstream from the anastomosis, with an average value of 3 Pa across the 4 mm fistula region. By day 21, peak TAWSS shifts to 1.2 mm downstream from the anastomosis, with the average value increasing fourfold to 13 Pa across the 4 mm fistula region.

By day 21, TAWSS showed significant changes in both magnitude and distribution. The peak TAWSS location shifted further downstream to 1.2 mm from the anastomosis, and its average value increased substantially to 13 Pa – a fourfold increase from day 7. This shift in peak location and magnitude suggests progressive vascular remodeling and potential stenosis development.

The three-dimensional TAWSS distribution (Fig 8) revealed further spatial variations. On day 7, peak TAWSS values were observed on the artery wall, with the maximum occurring at the inner side of the anastomosis on the proximal artery wall (Point A, 142 Pa) and a secondary peak at the stenosis (panel (a) Point B, 37 Pa). The proximal and distal arteries showed average TAWSS values of approximately 50 Pa and 4 Pa, respectively. By day 21, the TAWSS distribution pattern shifted significantly due to severe stenosis development in the fistula. The maximum TAWSS relocated to the AVF wall (panel (b) Point B, 200 Pa), with a secondary peak at the first stenosis (Point A, 134 Pa). The proximal and distal artery TAWSS values increased moderately to 60 Pa and 5 Pa, respectively.

**Fig 8.**
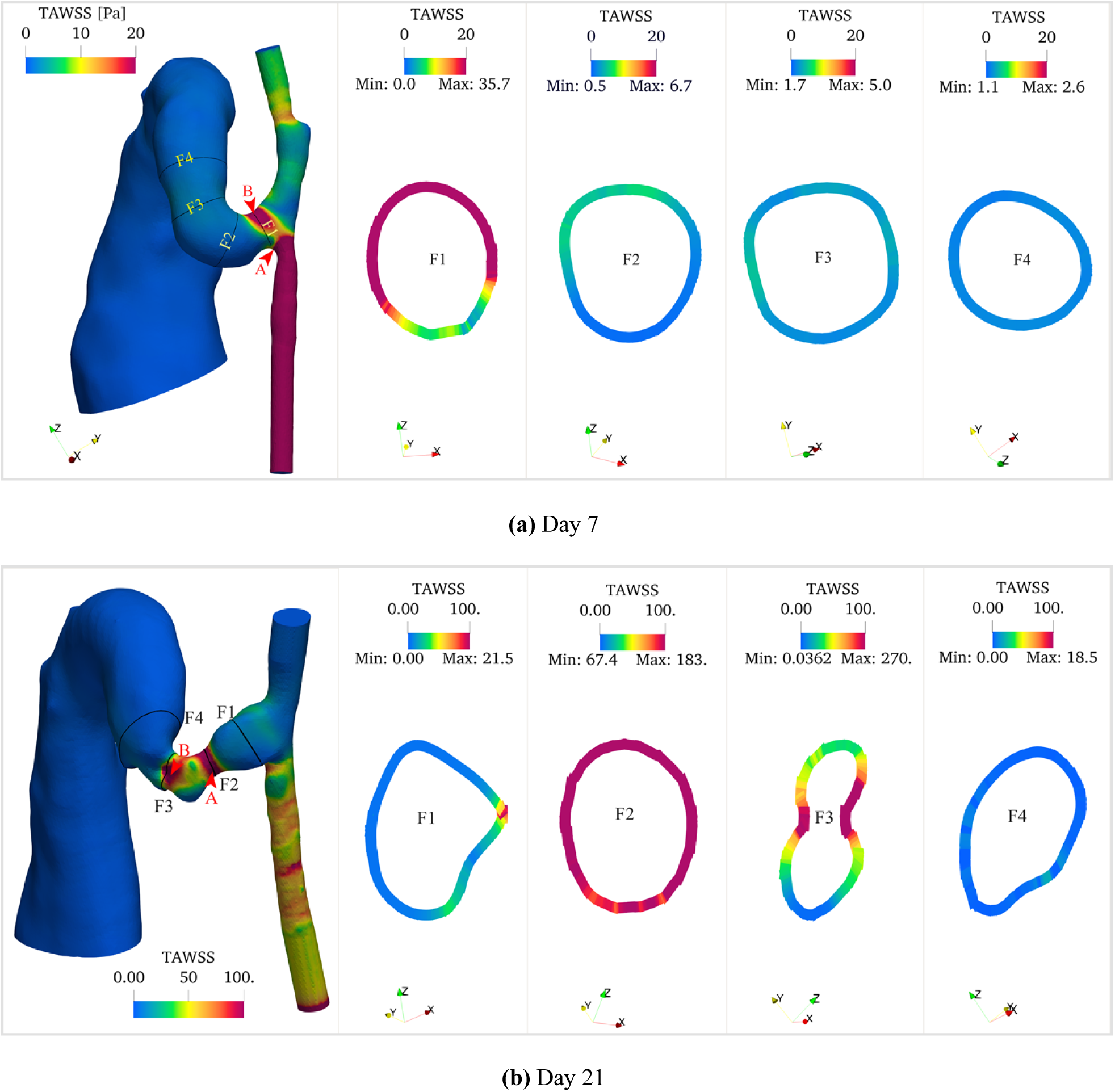
Time-averaged wall shear stress (TAWSS) distributions of (a) day 7 and (b) day 21 models. Each panel shows three-dimensional TAWSS colormaps along the vessel surfaces and representative cross-sectional (F1 to F4) distributions showing circumferential variation of TAWSS at key locations (anastomosis, stenosis, and AVF). TAWSS exhibits pronounced spatial heterogeneity, with peak values localized at the stenotic region. A significant temporal increase in TAWSS magnitude is observed from day 7 to day 21, particularly in regions of geometric constriction. Color scale represents TAWSS in Pa. Points A and B (indicated by arrows) show locations of maximum TAWSS values: 142 Pa and 37 Pa on day 7, and 134 Pa and 200 Pa on day 21, respectively. Cross-sectional dimensions are not to scale.

#### Oscillatory shear index analysis

The centerline analysis of OSI (Fig 9) reveals distinct temporal evolution patterns. On day 7, the maximum OSI occurred within 1.0 mm downstream from the anastomosis, with a relatively low average value of 0.0025 across the 4 mm fistula region, indicating minimal flow oscillation at the early stage.

**Fig 9.**
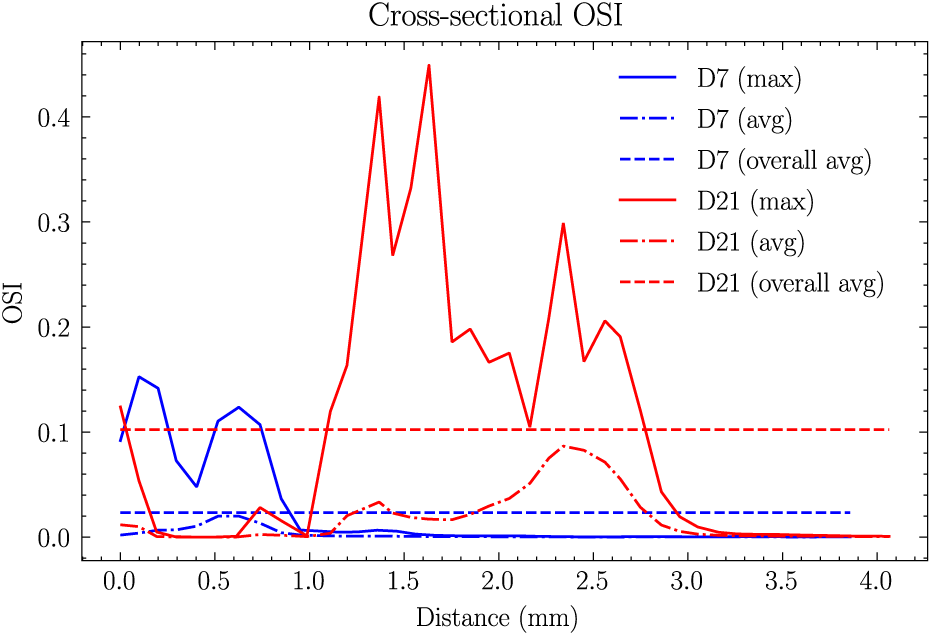
Cross-sectional maximum and average distributions of OSI along the centerline for (a) day 7 and (b) day 21. The two horizontal lines represent the average values across the first 4mm AVF. For day 7, peak OSI occurs within 1.0 mm downstream from the anastomosis, with an average value of 0.0025 across the 4 mm fistula region. By day 21, peak OSI extends from 1.0 to 2.5 mm downstream from the anastomosis, with the average value increasing significantly to 0.016 across the 4 mm fistula region.

By day 21, the OSI distribution showed extensive changes, with elevated values spanning a broader region from 1.0 to 2.5 mm downstream from the anastomosis. The average OSI increases more than sixfold to 0.016 across the 4 mm fistula region, indicating the development of more complex, disturbed flow patterns.

The three-dimensional OSI distribution (Fig 10) showed that OSI remained consistently low along the artery wall except near the anastomosis region. High OSI regions were primarily concentrated in two areas: at the anastomosis and downstream of stenotic segments, showing an inverse spatial relationship with TAWSS. The circumferential OSI distribution showed marked heterogeneity, particularly in regions of flow separation, indicating asymmetric flow behavior. Temporal analysis revealed a substantial increase in OSI magnitude from day 7 to day 21, most notably at the outside of the anastomosis (Point A, increasing from 0.22 to 0.48) and at Point B (increasing from 0.17 to 0.26).

**Fig 10.**
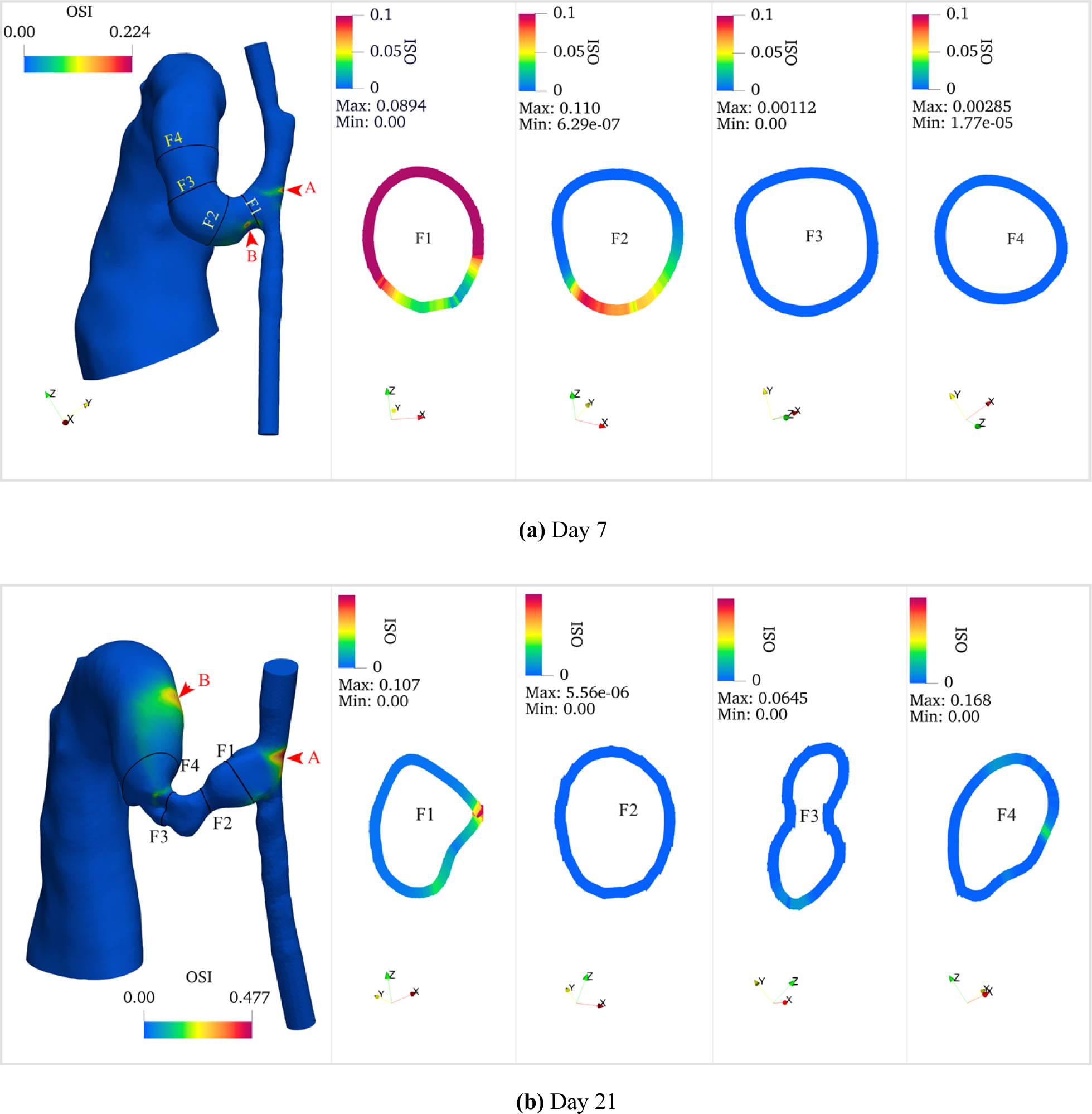
Oscillatory Shear Index (OSI) distributions of (a) day 7 and (b) day 21 models. Each panel shows three-dimensional OSI colormaps along the vessel surfaces and representative cross-sectional distributions showing circumferential variation of OSI at key locations (anastomosis, stenosis, and AVF). High OSI regions are predominantly localized at the anastomosis and downstream of the stenosis. OSI demonstrates circumferential heterogeneity, with elevated values corresponding to regions of flow separation. An obvious temporal increase in OSI magnitude is observed from day 7 to day 21. Color scale represents dimensionless OSI values from 0 to 0.1. Points A and B (indicated by arrows) show locations of maximum OSI values: 0.22 and 0.17 on day 7, increasing to 0.48 and 0.26 on day 21, respectively. Cross-sectional dimensions are not to scale.

#### Vortical structure

The development and evolution of vortical structures were analyzed using Q-criterion (Fig 11). At both timepoints, vortical structures concentrated predominantly near vessel walls, especially in regions of geometric transition, such as the anastomosis and stenotic segments. These wall-adjacent vortices developed primarily because of flow separation and adverse pressure gradients associated with the complex geometry. Notably, a distinct vortex core formed within the AVF vein lumen, indicating significant secondary flow patterns.

**Fig 11.**
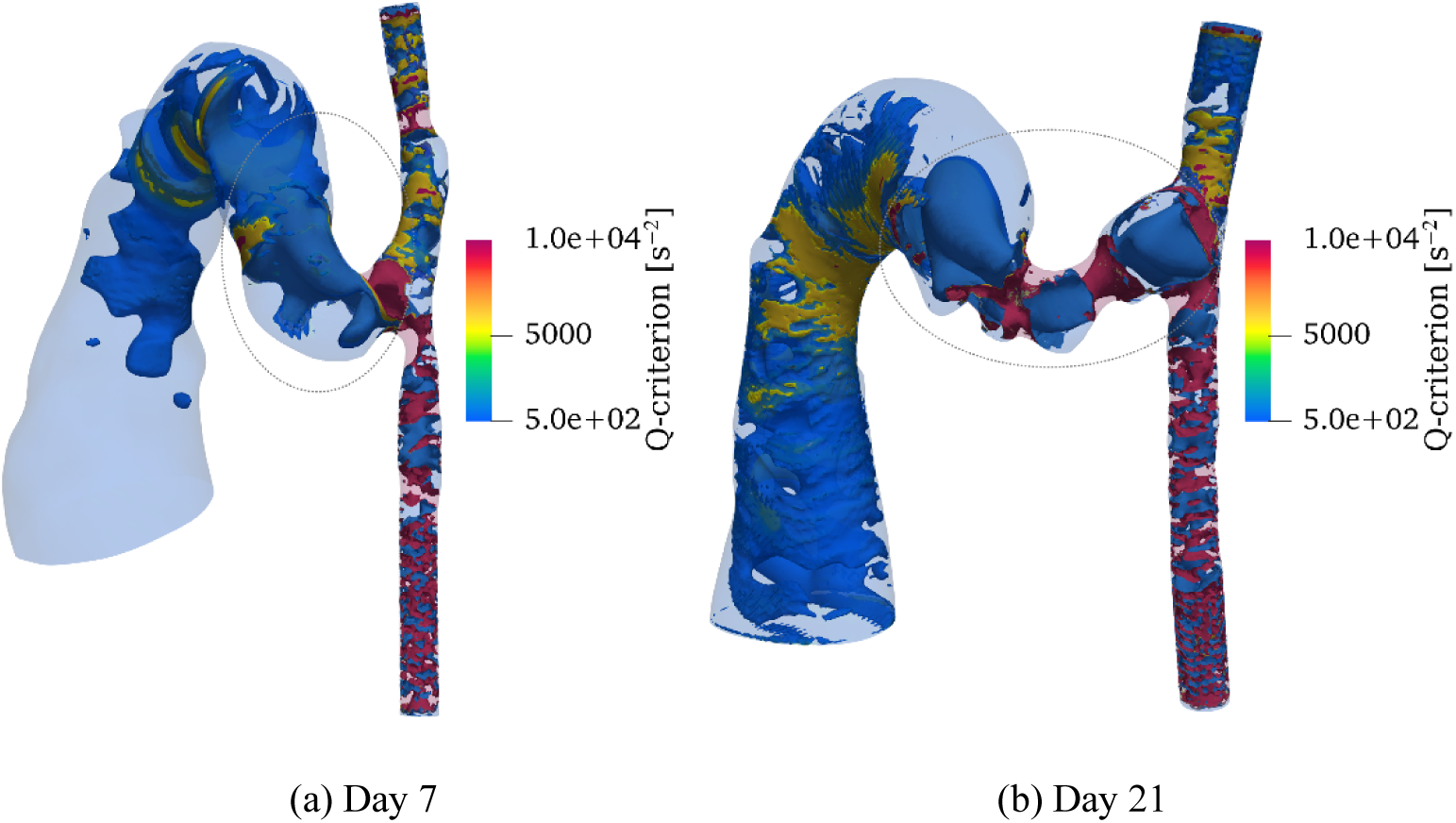
Vortical structures visualized using Q-criterion for (a) day 7 and (b) day 21 models. While vortical structures predominantly develop near vessel walls, a distinct vortex core (marked by circle) forms within the AVF vein lumen. Temporal comparison reveals increased vortical intensity on day 21 compared to day 7, particularly pronounced in stenotic regions. Although relatively small in magnitude, vortical structures are also observed to develop within the main vein. Color scale represents Q-criterion magnitude (*s*^−2^), with positive values indicating regions where rotation dominates strain.

The intensity and spatial distribution of vortical structures showed marked changes by day 21. The most noteworthy intensification occurred in stenotic regions, where the combination of flow acceleration and deceleration generated stronger rotational flow patterns. This enhanced vortical activity suggests increased flow complexity and higher energy dissipation in these regions. While vortical structures were also present in the main vein, their magnitude remained modest compared to those in stenotic regions and near-wall areas.

### Correlation analysis

The relationship between vessel geometry and hemodynamic parameters was analyzed through same-day correlations. Same-day correlations (Fig 12) showed a strong inverse relationship between TAWSS and vessel diameter, following fluid dynamic principles where smaller diameters exhibited higher wall shear stress under constant flow.

**Fig 12.**
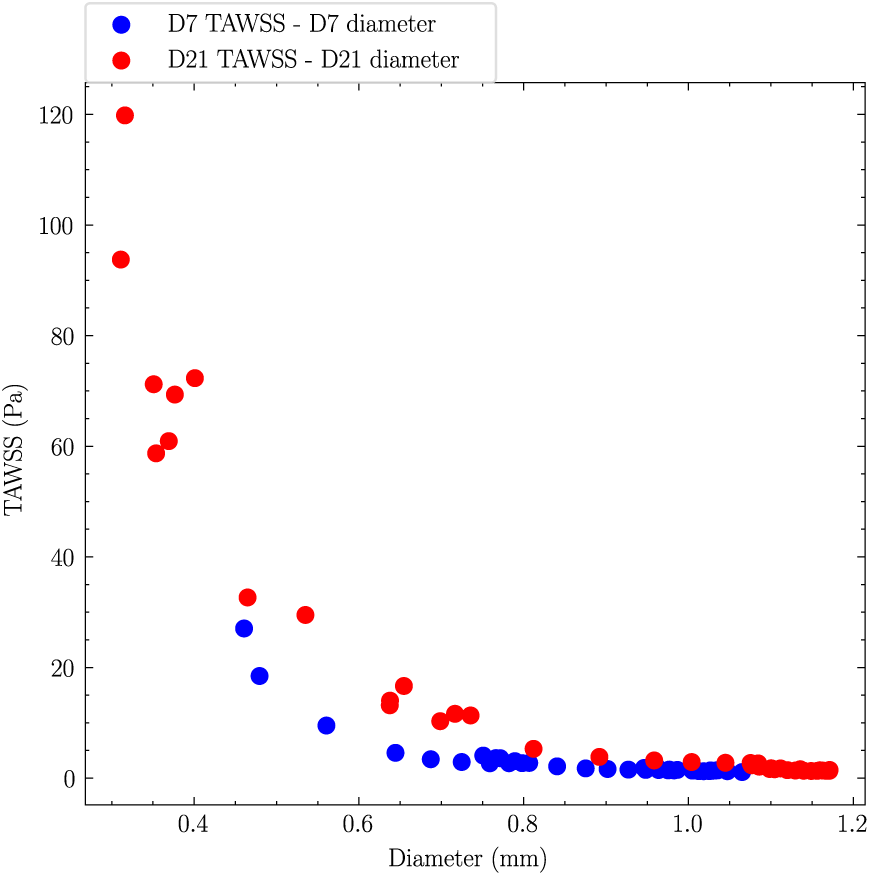
Same-day correlations between TAWSS and AVF lumen diameter. A strong nonlinear inverse relationship with diameter on both days was observed.

An inverse relationship between TAWSS and OSI was observed (not shown), consistent with previous findings in various vascular studies [36, 41], supporting the validity of our model.

Although the limited sample size prevented comprehensive statistical analysis of correlations between hemodynamic parameters and geometric changes, this study demonstrates the technical feasibility of identifying early hemodynamic conditions that predict subsequent remodeling and stenosis development.

The methodology established here can be applied to larger cohorts to identify associations between early hemodynamic parameters (TAWSS and OSI) and subsequent diameter changes in specific regions. This approach enables analysis of maximum and minimum stenosis at later stages (day 21) in relation to the TAWSS and OSI values at the same locations on day 7, potentially providing predictive markers for AVF maturation outcomes.

### Limitations and future directions

Several limitations of the present study should be acknowledged. First, our CFD model employed a simplified approach that assumed rigid vessel walls. While this assumption allowed for efficient computational analysis, it may not fully capture the vessel wall dynamics such as pulsatile deformation and wall vibrations that occur in vivo. Second, while our study demonstrated the feasibility of using CFD models to analyze hemodynamic correlations in AVF development, the small sample size limits the statistical power of our findings.

Future studies should address these limitations through two key improvements. Fluid-structure interaction simulations will better capture the dynamic interaction between blood flow and vessel wall mechanics. Additionally, an increased sample size will enable statistical validation of the observed correlations and potentially reveal new relationships between hemodynamic parameters and vascular remodeling, particularly in using early AVF CFD parameters to predict the progression of stenosis development and AVF maturation.

## Conclusion

This study demonstrates the utility of micro-CT-based computational fluid dynamics in characterizing the hemodynamic environment of murine AVF. Our analysis revealed significant temporal evolution in both geometric and hemodynamic parameters from day 7 to day 21 post-surgery. The AVF flow rate nearly doubled, while the fistula exhibited complex remodeling patterns with regional variations in diameter changes. The Wall shear stress analysis showed a shift in peak TAWSS locations and magnitude, accompanied by increased OSI, particularly at the region of anastomosis.

The methodology developed in this study enables analysis of relationships between early hemodynamic conditions and subsequent geometric changes, establishing a framework for identifying potential predictive indicators of stenosis development. These insights into the mechanistic relationship between hemodynamics and vascular remodeling may also be helpful to develop strategies to improve AVF outcomes in clinical practice, by changing the anastomotic angle in order to achieve future optimized flow profiles that would result in improved AVF maturation. Through the combination of high-resolution micro-CT imaging with ultrasound flow data, our approach not only provides a framework for future investigations into vascular adaptation and dysfunction in AVF development but also carries the potential for an easy and affordable predictive test that can inform the type (angle and curvature) of created AVF in the setting of a particular baseline artery-vein configuration.

## Acknowledgements

G.X. and P.R.C. acknowledge funding from the NIH (Award R01DK132328). E.M.B acknowledges funding from the NIH (Award R56DK140967). B.E.G. acknowledges funding from the NIH (Awards R01HL157631 and U01HL143336) and NSF (Award OAC 1931516).

Simulations were performed using computational facilities provided by the University of North Carolina at Chapel Hill through the Research Computing Division of UNC Information Technology Services.

